# Correlative multiscale 3D imaging of mouse primary and metastatic tumors by sequential light sheet and confocal fluorescence microscopy

**DOI:** 10.1101/2024.05.14.594162

**Authors:** Jingtian Zheng, Yi-Chien Wu, Xiaoying Cai, Philana Phan, Ekrem Emrah Er, Zongmin Zhao, Steve Seung-Young Lee

## Abstract

Three-dimensional (3D) optical microscopy, combined with advanced tissue clearing, permits *in situ* interrogation of the tumor microenvironment (TME) in large volumetric tumors for preclinical cancer research. Light sheet (also known as ultramicroscopy) and confocal fluorescence microscopy are often used to achieve macroscopic and microscopic 3D images of optically cleared tumor tissues, respectively. Although each technique offers distinct fields of view (FOVs) and spatial resolution, the combination of these two optical microscopy techniques to obtain correlative multiscale 3D images from the same tumor tissues has not yet been explored. To establish correlative multiscale 3D optical microscopy, we developed a method for optically marking defined regions of interest (ROIs) within a cleared mouse tumor by employing a UV light-activated visible dye and Z-axis position-selective UV irradiation in a light sheet microscope system. By integrating this method with subsequent tissue processing, including physical ROI marking, reversal of tissue clearing, tissue macrosectioning, and multiplex immunofluorescence, we established a workflow that enables the tracking and 3D imaging of ROIs within tumor tissues through sequential light sheet and confocal fluorescence microscopy. This approach allowed for quantitative 3D spatial analysis of the immune response in the TME of a mouse mammary tumor following cancer immunotherapy at multiple spatial scales. The workflow also facilitated the direct localization of a metastatic lesion within a whole mouse brain. These results demonstrate that our ROI tracking method and its associated workflow offer a novel approach for correlative multiscale 3D optical microscopy, with the potential to provide new insights into tumor heterogeneity, metastasis, and response to therapy at various spatial levels.

## Introduction

A solid tumor presents a complex structure at various scales^1,2^. At the whole tumor scale, clinical imaging and surgical operations have revealed tumors with various shapes and sizes in cancer patients^3,4^. At the tissue scale, numerous abnormal features, such as neoplasia, desmoplasia, hypoxic and necrotic areas, random immune infiltrates, and haphazard vasculature, have been identified in the tumor microenvironment (TME)^5–7^. At the cellular scale, a variety of cell types and their intricate organization in different subregions of a tumor have recently been unveiled^8,9^. Each of these features, across various scales, significantly affects tumor progression and treatment outcomes^10–12^. Therefore, studying tumor architecture at these scales is important for a better understanding of tumor development and for developing effective therapies.

Recent advances in tissue clearing-based 3D optical microscopy have offered unprecedented 3D images of tumor tissues at multiple resolutions^13,14^. Simple immersion in a clearing solution with a high refractive index (RI, ∼1.44-1.56) allows sufficient transparency in tumor tissues for 3D optical microscopy^15^. For 3D light sheet fluorescence microscopy (LSFM, or ultramicroscopy), organic solvents such as dibenzyl ether (DBE) and ethyl cinnamate (ECi) have been widely used for optical clearing of whole mouse tumor tissues after dehydration in alcohol (e.g. ethanol (EtOH))^16,17^. LSFM offers macroscopic and mesoscopic resolution 3D images of the cleared mouse tumor tissues using objectives with a low-magnification (1x or 4x) and long working distance^18,19^. In contrast, confocal laser scanning microscopy (CLSM) has generally been utilized for imaging tumor tissues at cellular resolution^20,21^. By integrating with aqueous sugar solution (e.g., D-fructose)-based tissue clearing and multiplex immunofluorescence (IF) methods, CLSM can produce high-resolution 3D images of multiple cell types throughout a half millimeter-thick tumor tissue^22,23^. LSFM and CLSM provide distinct spatial resolutions and fields of view (FOVs) in the 3D imaging of tumor tissues. Notably, our previous work demonstrated that switching the clearing solution from an organic solvent to an aqueous sugar solution permits the sequential LSFM and CLSM of the same mouse tissue sample^24^. The sequentially combined microscopy process can produce comprehensive spatial information across whole mouse organ, tissue, and cellular levels. However, after LSFM, vibratome-mediated macrosectioning is required for high-resolution 3D CLSM of the cross-sectioned area of the tissue sample. This physical tissue macrosectioning poses challenges in tracking regions of interest (ROIs) and in registering collected multiscale 3D images.

To address this challenge, we have developed a novel method that visibly labels the ROI positions in tissues. We infused agarose gel surrounding the cleared tissue with UV-activated (UVA) dye molecules, spiropyran, and then exposed these ROI positions to a UV light sheet beam in the light sheet microscope system for optical marking. The downstream tissue processes, which involve physical ROI marking, switching clearing solutions, vibratome macrosectioning, and IF, facilitate the acquisition of tissue macrosections that retain the ROIs for high-resolution multiplex CLSM. Our developed workflow has ultimately enabled us to identify, preserve and track ROIs within mouse tumor and brain tissues between 3D LSFM and CLSM imaging sessions. We validated the accuracy of our workflow in the acquisition of ROI-retained macrosections by embedding a single fluorescent microbead in an agarose gel plug as an artificial ROI. We further applied the workflow for evaluating the anti-cancer immune response in a TUBO mouse mammary tumor after treatment with a stimulator of interferon genes (STING) agonist, 5,6-dimethylxanthenone-4-acetic acid (DMXAA). At the whole tumor, tissue, and cellular scales, the multiresolution 3D tumor images revealed distinct spatial features of the TMEs in treated and untreated mouse tumors, while also being readily registered together to produce a correlated 3D image. In the tumor treated with DMXAA, quantitative 3D spatial analysis of LSFM images showed that intratumoral ROIs with intensified autofluorescence (excitation: 639 nm, emission: 665-695 nm) correspond to tissue areas exhibiting larger numbers of CD45+ immune cells and fewer CK8+ cancer cells, as confirmed by multiplex CLSM. In addition, our workflow enabled the detection and 3D multiplex imaging of a metastatic lesion within a whole brain tissue from a breast cancer metastasis mouse model. This work demonstrates that our ROI tracking method and sequential 3D LSFM-CLSM microscopy workflow present a novel approach for correlative multiscale 3D imaging of mouse primary and metastatic tumors.

## Results

### Workflow for correlative 3D LSFM and CLSM of tumor tissues

We established a robust workflow that incorporates a novel UVA dye-employed ROI tracking method for correlative ‘whole mouse tumor (or organ)-to-cell resolution’ 3D imaging *via* sequential LSFM and CLSM (**Figure 1** and **Table 1**). In this workflow, whole mouse tumor tissue (or mouse brain) is fixed in paraformaldehyde (PFA) solution, embedded in agarose gel, and optically cleared using an organic solvent-based method involving serial immersions in EtOH (20%, 50%, 100% v/v), DBE, and ECi solvents (**Supplementary Figure 1**). Adjusting the pH of these solvents to approximately 7.8 enhances the preservation of protein antigens in the tissue for downstream IF labeling of various cell markers^25,26^. Next, the cleared tissue undergoes 3D LSFM imaging to capture autofluorescence signal across the entire tissue sample using a 4× immersion objective, and 639 nm excitation laser and its corresponding emission filter (665-695 nm). The long-wavelength excitation laser beam offers superior tissue penetration^27,28^, enabling the scanning of autofluorescence originating from large volumetric mouse tumors and whole mouse brains by LSFM. The collected tile image data are reconstructed and rendered in 3D using Imaris software. Using interactive image view functions in the software, such as virtual 2D and orthogonal slicers, regions of interest (ROIs) within the 3D tissue image are defined. Afterwards, the tissue embedded in an agarose plug is pulled out from the sample chamber of the light sheet microscope, and then incubated in ECi solvent containing spiropyran to infuse the UVA dye into the agarose gel. Likely, adjusting the pH of the ECi solvent to 7.8 is necessary to activate the spiropyran in the gel upon exposure to a UV light sheet beam^29^. The tissue is then returned to the LSFM chamber, and the focus plane of light sheet beam is positioned at the height of the tissue presenting the ROI identified by the live scanning of tissue autofluorescence with 639 nm excitation. The light source is switched to a 405 nm UV laser, and a one-sided light sheet beam illuminates to activate the UVA dye molecules in the selected area of the gel plug at the ROI position. This enables optical marking of the ROI by generating a thin, purple-colored horizontal line in the agarose gel. After this optical ROI marking, the tissue is immediately pulled out from the LSFM chamber, and the surface of the agarose plug is cut at the purple-colored line using a razor blade. This physical marking of the ROI is required due to the reversible chemical transformation of spiropyran under visible light, which causes the purple-colored line to disappear^30^. Subsequently, the tissue undergoes rehydration and a reverse process of organic solvent-based tissue clearing. Guided by the physical mark showing the cut one-side edge of the gel, 400 µm-thick tissue macrosection retaining the ROI is obtained through vibratome sectioning. The tissue macrosection is labeled with fluorescent primary antibodies to localize various cell types. After optical clearing in D-fructose solutions, the detailed tissue-level structure and cellular composition of the whole tissue macrosection is visualized through 3D mosaic imaging by CLSM using a low-magnification objective (10× or 20×). Cellular resolution 3D images of the ROI within the macrosection can also be achieved by CLSM with a high-magnification objective (40× or 60×). The image data from LSFM and CLSM data are transferred to a PC workstation and registered into a correlative multiscale 3D image. As a result, a ‘Google Earth-like’ view of ROI tracking in the tissue sample and quantitative 3D spatial analyses at various scales are feasible.

**Figure 1.**
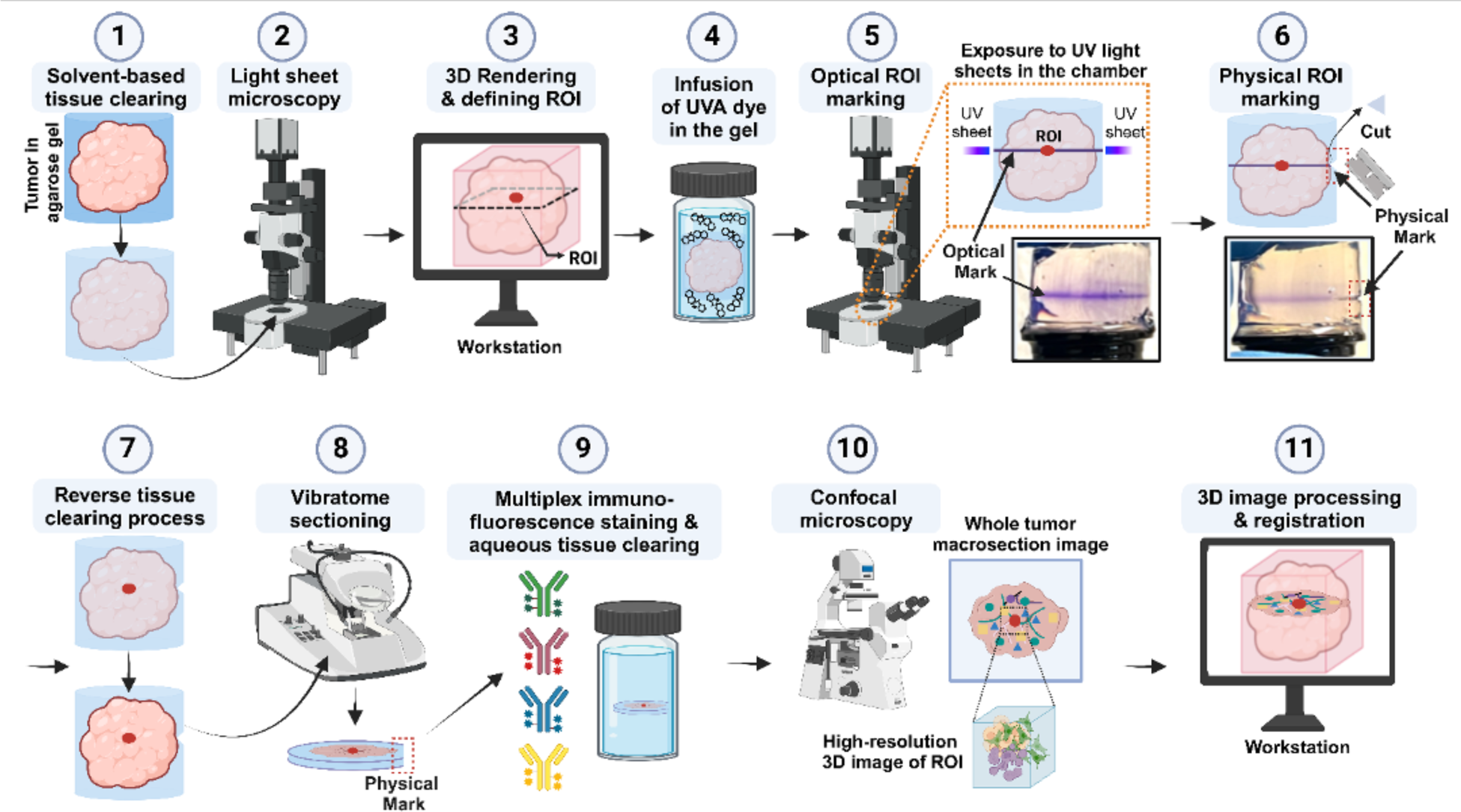
Workflow for correlative 3D LSFM and CLSM imaging of mouse tumors. **(1)** Optical clearing of a mouse tumor embedded in agarose gel by sequential incubation in EtOH, DBE, and ECi. **(2)** 3D LSFM scanning of autofluorescence in the whole tumor. **(3)** Defining ROI based on 3D visualization of reconstructed tumor image. **(4)** Infusion of UVA dye, spiropyran, into the agarose gel surrounding the tumor. **(5)** Optical marking of the ROI *via* exposure to UV light sheet beam in LSFM. The photograph (right bottom) shows a purple-colored line on an agarose plug as an optical mark. **(6)** Physical marking of the ROI by cutting the surface of the agarose gel along the purple-colored line. The photograph (bottom) displays a physical mark on the agarose gel, made using a razor blade. **(7)** Reversing the process of organic solvent-based tissue clearing. **(8)** Vibratome sectioning of the tumor with the guidance of the physical mark. The 400 µm-thick tumor macrosection (bottom), which retains the ROI, shows a one-sided loss of the agarose gel surrounding the tumor, serving as a physical mark. **(9)** Multiplex immunofluorescence staining and optical tissue clearing of the tumor macrosection in D-fructose solution. **(10)** 3D multiplex CLSM imaging of the tumor macrosection at both tissue-wide and cellular resolution. **(11)** 3D image processing, registration, and quantitative analysis.

**Table 1.** Overview of the procedure and average times required for each step.

| Step | Process | Average Time | Notes |
| --- | --- | --- | --- |
| 1 | Organic solvent-based tissue clearing | 5-7 days | Harvest tumor and other tissues<br>Embed the tissue in 2% agarose gel<br>Dehydration in 20, 50, and 100% EtOH<br>Optical clearing in DBE and ECI |
| 2 | Light sheet fluorescence microscopy (LSFM) | Hours (depends on the size of tissue) | Place the cleared tissue in the chamber of LSFM filled with ECI solvent<br>Record the X-Y orientation of the sample on the stage<br>Image the whole tissue by LSFM with a specific excitation and the corresponding emission filter |
| 3 | 3D rendering & defining ROI | Varies | Export image data and 3D rendering in Imaris software<br>Define ROI by screening virtual cross-sections |
| 4 | Infusion of UVA dye in the gel surrounding the tissue | ~12 hrs | Incubate the sample in spiropyran dissolved in ECI for overnight under protection from light at room temperature |
| 5 | Optical marking of ROI | ~30 mins (3 mins for exposing to UV light sheet beam) | Placed the tissue in the chamber of LSFM<br>Search for the defined ROI by scanning along Z-axis with the same laser setting used in the first imaging<br>Expose the tissue and gel to a one-sided UV light sheet beam at the Z-axis position corresponding to the ROI |
| 6 | Physical marking of ROI | 5 secs | Take out the tissue, quickly cut the edge of the agarose gel along the optically marked 'purple' colored line |
| 7 | Reverse tissue clearing process | 3 days | Rehydrate the tissue by incubation in 100, 50, and 20% EtOH and PBS in sequence |
| 8 | Vibratome sectioning | ~30 mins | Section the tissue at 400 μm thickness in serial and identify the macrosection with a cut gel edge. |
| 9 | Immunofluorescence staining, & aqueous tissue clearing | 2 days | Stain the macrosection with fluorescent antibodies cocktail overnight<br>Incubate in 20, 50, 80, and 100% (w/v) D-fructose solution in sequence, each for 30 mins |
| 10 | Confocal laser scanning microscopy (CLSM) | Varies | Image the whole tissue macrosection and ROI by multiplex 3D CLSM using 20x and 40x objectives |
| 11 | 3D image processing & registration | Varies | Construct the correlated multiscale 3D image using Fiji and Imaris software<br>Segment and analyze the image data using Fiji software |

To validate the precision cutting of an ROI-inclusive tissue macrosection, a single fluorescent microbead, 180-200 µm in diameter, was placed in the middle of an agarose gel plug to create an artificial ROI (**Supplementary Figure 2**). At the end of the workflow, the gel plug was subjected to the organic solvent-based tissue cleaning process. LSFM identified the location of the microbead within the plug. The optical and physical markings of the ROI on the surface of the gel plug were made according to the protocols using spiropyran. The gel plug was serially macrosectioned to a thickness of 400 µm using a vibratome. The gel macrosection retaining the microbead was readily identified by finding the physical mark on the edge of the gel. CLSM imaging confirmed that the microbead was within the selected macrosection. This result demonstrates that ROI tracking mediated by UVA dye allows for the direct identification of tissue macrosections retaining ROIs for correlative multiscale 3D imaging.

### Tracking of ROIs in mouse tumors treated with immunotherapy

To model the imaging assay of immunotherapy effects on human breast tumors, we applied our workflow to mouse mammary TUBO tumors, both with and without the intratumoral injection of DMXAA. After collecting tumors, embedding in agarose gel, and optical clearing with organic solvents, both control and treated tumor tissues underwent LSFM imaging using a 639 nm excitation laser and related emission filter. The 3D images of whole tumor tissues were reconstructed based on their tissue autofluorescence signals using Imaris software.

Using 3D rendering and virtual 2D cross-section display functions, two horizontal planes containing ROIs in each tumor were selected (**Figure 2A**). There was a relatively even distribution of autofluorescence signals across the selected planes in the control tumor. In contrast, varying signal intensities were observed over the virtual cross-section images of the treated tumor (**Figure 2B**). We selected areas expressing low- and high-intensity autofluorescence signals as the ROIs in each virtual plane of the treated tumor. By applying optical and physical ROI marking methods, two 400 µm-thick tumor macrosections containing the ROIs were selectively obtained from each control and treated tumor (**Figure 2C**). After optically clearing the tumor macrosections in D-fructose solutions, we imaged them using CLSM with a 640 nm excitation laser and its matched emission filter. Subsequently, we reconstructed the 3D images of the entire macrosections based on their autofluorescence signals, which were captured using similar excitation and emission settings as in LSFM. Using the virtual 2D cross-section viewing mode in Imaris, we identified the individual ROI-containing planes in the middle of the 3D tumor macrosection images (**Figure 2D**). For quantitative measurement of the accuracy of our ROI tracking method, the similarities between the virtual tumor cross-section images acquired by LSFM and CLSM were assessed using the structural similarity index measure (SSIM) method (**Figure 3**)^31^. SSIM ratios for the distribution patterns of tissue autofluorescence signals across the whole virtual cross-section images were calculated after converting the image data to 8-bit binary form and removing non-specific background signals outside the tumor area (**Figure 3A**). The outer layers of the tumors in the binary images were also segmented to separately determine similarity indexes for the patterns of tumor outlines in the virtual cross-section images from LSFM and CLSM. The yellow areas in the merged images represent overlapping patterns of autofluorescence signals and tumor outlines between LSFM (red) and CLSM (green) images. The average SSIM ratios for the whole cross-section and the outer layer of the tumor images from LSFM and CLSM were 0.68 and 0.89, respectively (**Figure 3B**). These SSIM ratios indicate a high level of similarity between the LSFM and CLSM images. For comparison, we used identical LSFM and CLSM images to determine an SSIM ratio of 1 for the positive control. Additionally, we cross-compared LSFM and CLSM images from different tumor groups to calculate an average SSIM ratio of 0.15 for the negative control. These results demonstrate the high accuracy of our method in tracking ROIs within tissue samples throughout sequential LSFM and CLSM imaging processes.

**Figure 2.**
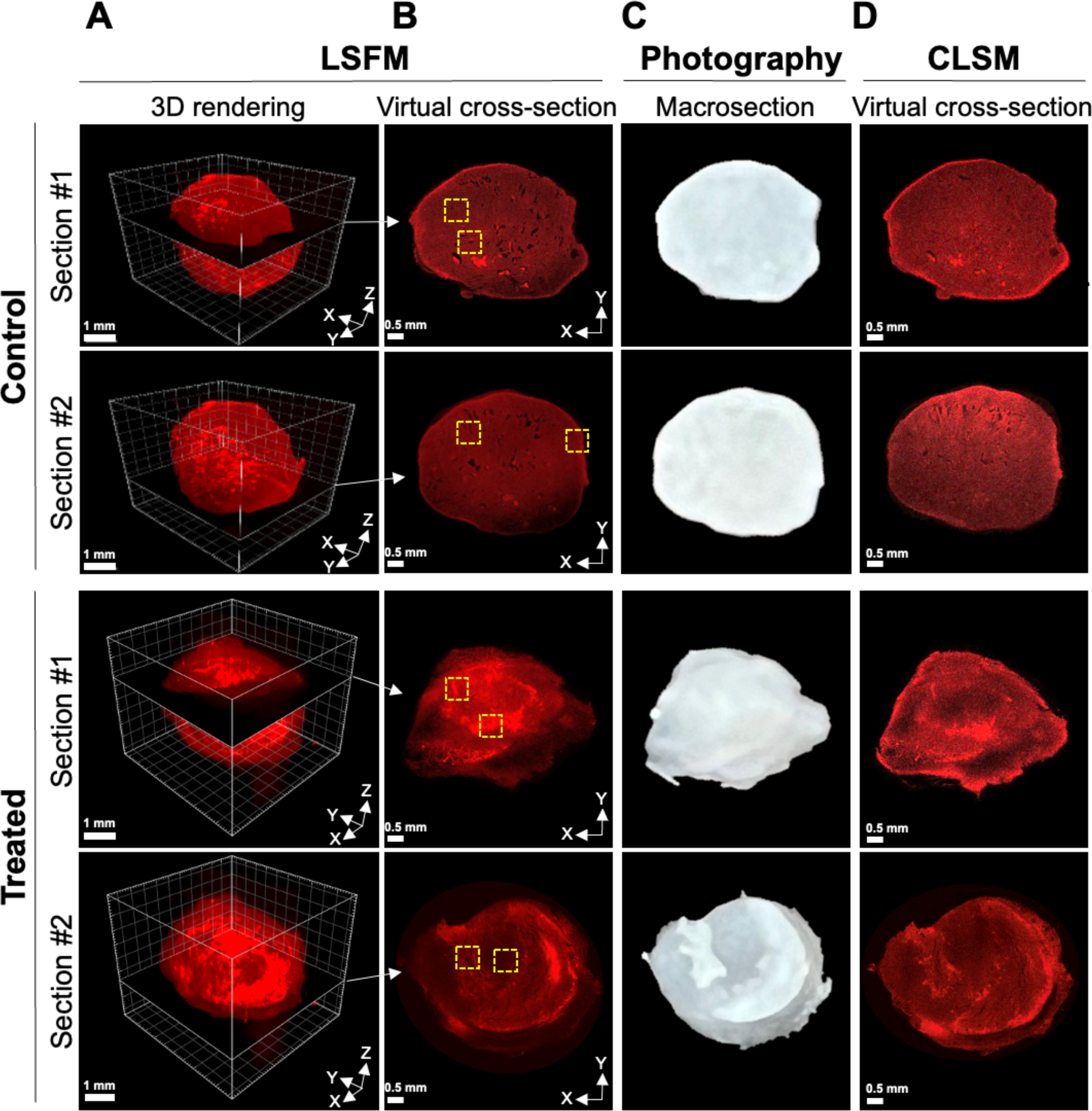
Tracking ROIs in the control and DMXAA-treated mouse tumors under sequential LSFM and CLSM imaging. **(A)** 3D rendering and virtual slicing of the whole control and treated tumor images acquired by 3D LSFM scanning of autofluorescence signals. Scale bar: 1 mm. **(B)** Virtual cross-section views containing ROIs. Yellow dotted boxes indicate ROIs in each virtual tumor cross-section. Scale bar: 0.5 mm. **(C)** Photographs of 400 µm-thick tumor macrosections retaining ROIs, which were obtained through optical and physical ROI marking and vibratome sectioning. **(D)** Virtual cross-section views of the macrosection images displaying the ROIs, acquired through 3D CLSM scanning of autofluorescence signals. Scale bar: 0.5 mm.

**Figure 3.**
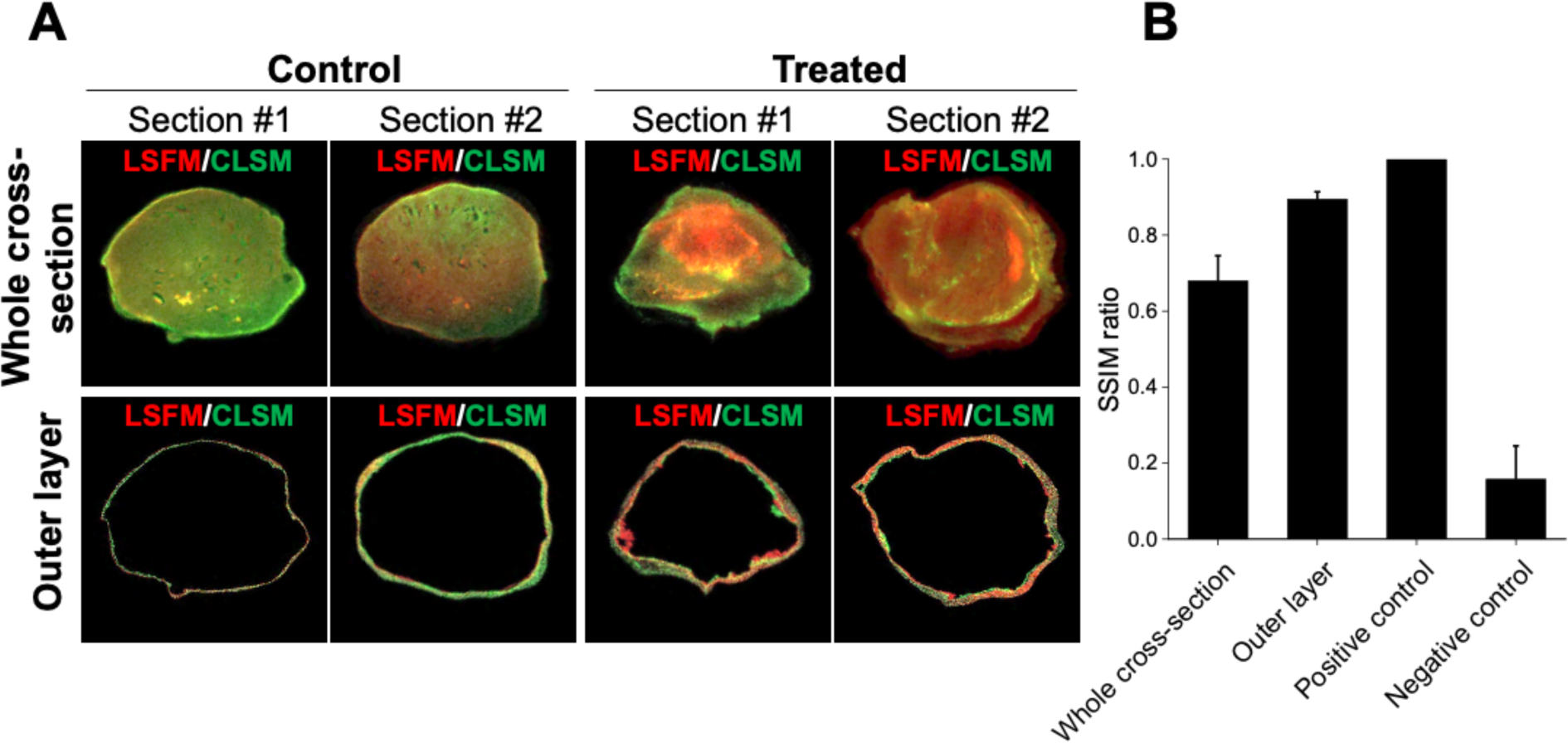
Quantitative measurement of similarity between the virtual tumor cross-section images from LSFM and CLSM. **(A)** Merged images of virtual whole tumor cross-sections (top row) and only their outer layers (bottom row). The virtual tumor cross-section images acquired by LSFM and CLSM were colored red and green, respectively. **(B)** SSIM ratios between LSFM and CLSM images in the forms of virtual whole tumor cross-section and outer layer (n=4, including section #1 and #2 in both control and treated groups). The SSIM ratio of the positive control was determined using identical LSFM or CLSM images. The SSIM ratio of the negative control was measured by cross-comparing LSFM and CLSM whole cross-section images between different tumor groups.

### Correlative multiscale 3D spatial analysis of cellular response to immunotherapy in the tumor microenvironment

To investigate the effects of DMXAA treatment on the spatial cellular composition of the TME, we collected macrosections containing ROIs from both control and treated tumors using our ROI tracking protocol. The macrosections were stained with DAPI, DyLight488-anti-CK8, DyLight550-CD45, and DyLight680-anti-ER-TR7 antibodies to localize cell nuclei, cancer cells, immune cells, and fibroblasts, respectively. After optical clearing in D-fructose solutions, we imaged the labeled macrosections using 3D multiplex fluorescence CLSM with 20× and 40× objectives to achieve tissue-wide and cellular resolution 3D images (**Figure 4**). The 3D images of the whole macrosections revealed significant structural remodeling of the broad TME in the treated tumor as compared to the control group (**Figure 4A**). A high density and even distribution of CK8+ cancer cells were observed throughout the control tumor macrosections. In contrast, the treated tumor macrosections displayed a low density, scattered distribution, and regional loss of CK8+ cancer cells. On the other hand, CD45+ immune cells were extensively distributed throughout the treated tumor macrosections, but were mostly localized to the edges of the control tumor macrosections.

**Figure 4.**
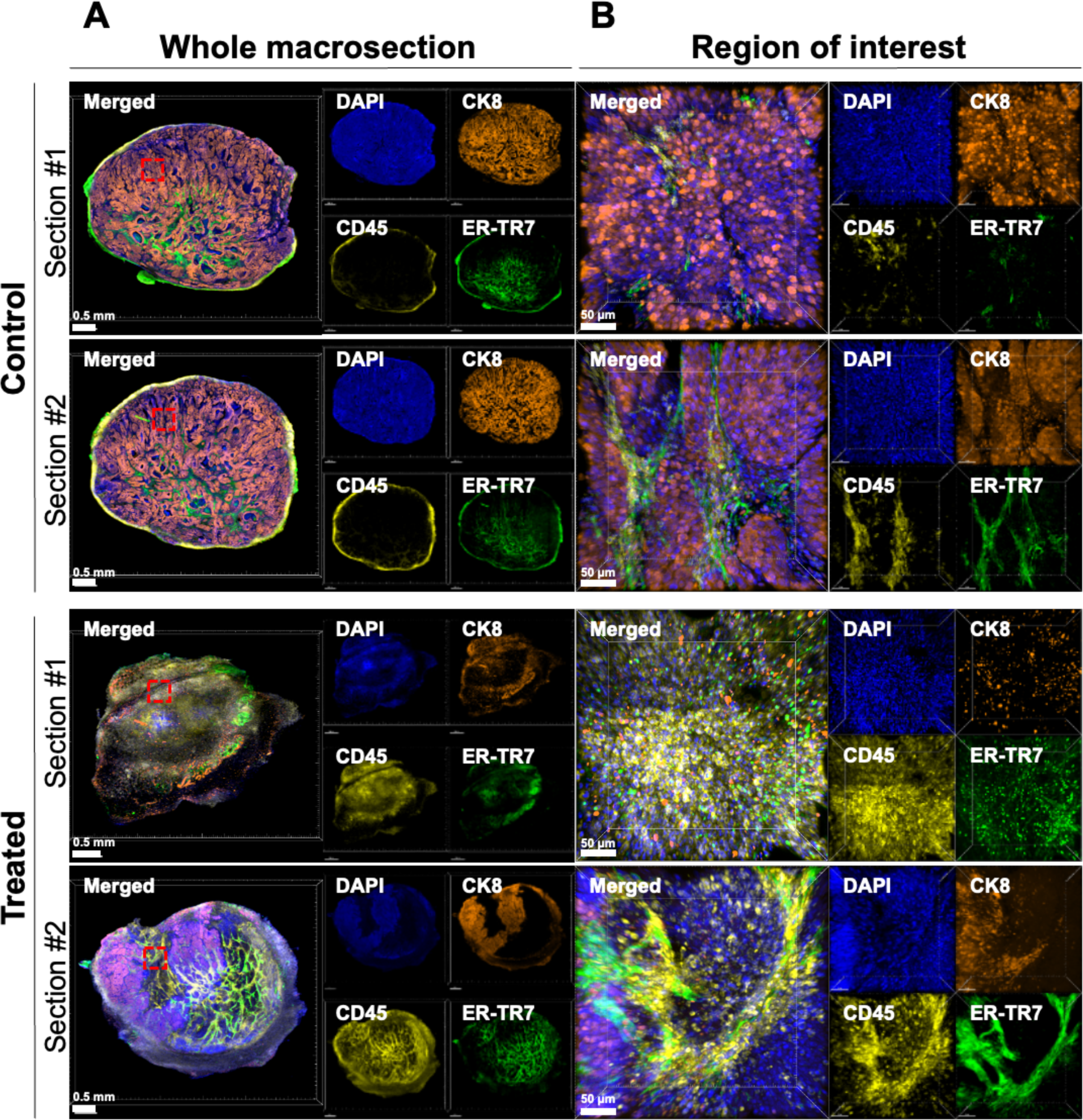
High-resolution 3D multiplex images of the whole tumor macrosections and defined ROIs in control and DMXAA-treated tumor groups. **(A)** 3D rendering of whole macrosection images from control and treated tumors acquired by CLSM. The 3D images display the locations of DAPI+ (blue) cell nuclei, CK8+ (orange) tumor, CD45+ (yellow) immune infiltrates, and ER-TR7+ (green) fibroblast-abundant stroma. Scale bar: 0.5 mm. (**B**) 3D rendering of ROI images obtained by high-resolution 3D CLSM of the tumor macrosections. High-resolution 3D images of other ROIs are in **Supplementary Figure 3**. Red dotted boxes within each tumor macrosection in **A** indicate selected ROIs. The cellular resolution 3D images exhibit the structural cell composition within the ROIs. Scale bar: 50 µm.

To quantify the differences, the areas positive for each cellular marker in the whole tumor macrosection images were extracted using hyperstack segmentation in Fiji software with optimal individual cutoff thresholds (**Figure 5A**). The total volume percentages (%) of CK8+, CD45+, and ER-TR7+ cells in each macrosection were calculated by V % = (Total volume of each cell type / Total volume of DAPI+ tumor parenchyma) × 100 (**Figure 5B**). In alignment with our observations, smaller volumes of CK8+ cancer cells were calculated in the macrosections of the treated tumor (#1: 10%, #2: 38%) compared to those in the control tumor (#1: 78%, #2: 79%). For the immune cell component, larger volumes of CD45+ immune cells were measured in the macrosections of the treated tumor (#1: 75%, #2: 38%) compared to those in the control tumor (#1: 9%, #2: 10%). Additionally, we examined the distribution patterns of CD45+ immune cells over the CK8+ tumor area in representative macrosections from each tumor group (**Figure 5C**). Intensity profiles of CK8+ (green) and CD45+ (red) signals along the white lines across the macrosections clearly demonstrated that most CD45+ immune cells were localized on the outside of the CK8+ tumor parenchyma in the control tumor macrosection, while CD45+ immune cells were distributed throughout the treated tumor macrosection, indicating significant immune infiltration in response to DMXAA treatment^32^.

**Figure 5.**
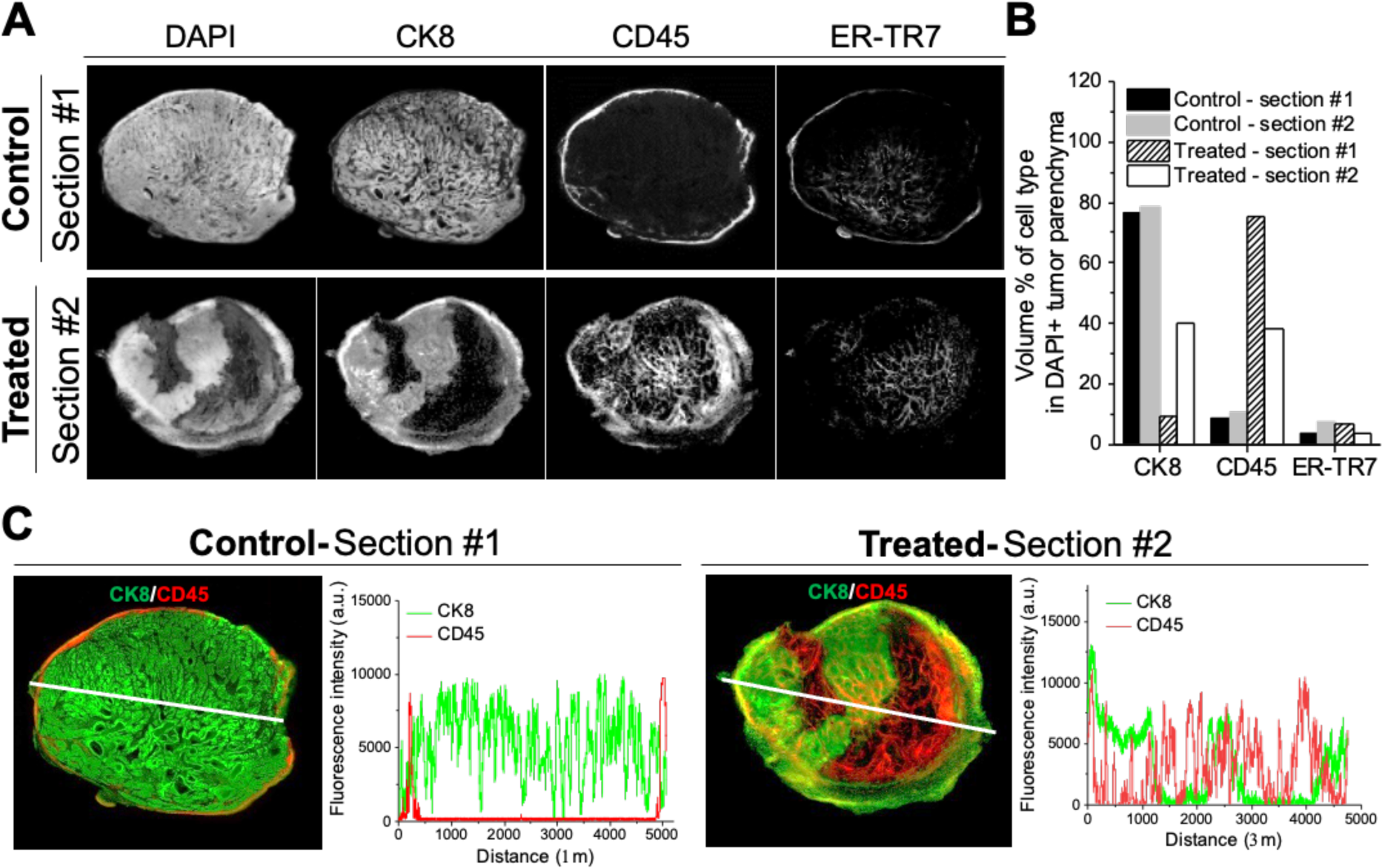
Quantitative 3D spatial analysis of cellular response to immunotherapy in the tumor microenvironment. **(A)** Segmentation of individual cell markers in representative control and treated tumor macrosection images (Section #1 in the control tumor, Section #2 in the treated tumor). **(B)** Volume percentages (%) of CK8+, CD45+, and ER-TR7+ segments normalized with DAPI+ parenchyma in individual control and treated tumor macrosections. **(C)** Distribution patterns of CK8+ tumor cells and CD45+ immune cells in the representative control and treated tumor macrosections. Merged 16-bit channel images for CK8+ (green) and CD45+ (red) in the tumor macrosections (**left**). Fluorescence intensity profiles of CK8+ (green) and CD45+ (red) signals along the lines indicated in the each macrosection image (**right**).

CLSM also provided cellular resolution 3D images of the ROIs within the macrosections using a high magnification objective (40×) (**Figure 4B and Supplementary Figure 3**). The high-resolution 3D images revealed distinct distribution patterns of CD45+ immune cells in the ROIs between the control and treated tumors. For quantitative 3D spatial analysis of immune infiltration at cellular resolution, the fluorescence signals of individual CK8+, CD45+, and ER-TR7+ cells in the ROI images were segmented using Imaris software. The segmented images of these different cell types were rendered in 3D (**Figure 6A**). We then calculated the volume percentages (%) of CK8+ cancer cells that interacted with CD45+ immune cells in the 3D space (**Figure 6B**). Approximately 35% of CK8+ cancer cells spatially contacted CD45+ immune cells in the ROIs of the control tumor, while in the treated tumor, around 80% of CK8+ cancer cells interacted with CD45+ immune cells. Additionally, we measured the 3D penetration distances of CD45+ immune cells from ER-TR7+ fibroblast-abundant stroma to tumor parenchyma (**Figure 6C**). The mean tumor-infiltrating distances of immune cells in the ROIs of the control and treated tumors were 5 µm and 14 µm, respectively. These findings suggest that the eradication of cancer cells was attributable to increased infiltration of immune cells into the tumor parenchyma following DMXAA treatment^33^. Interestingly, ROIs showing high immune infiltration, as visualized by multiplex CLSM imaging, correlated with areas exhibiting high autofluorescence signals in the virtual cross-sections of LSFM images. Conversely, ROIs with a high density of cancer cells in the multiplex CLSM images corresponded to areas of low autofluorescence signals in the virtual cross-sections of LSFM images.

**Figure 6.**
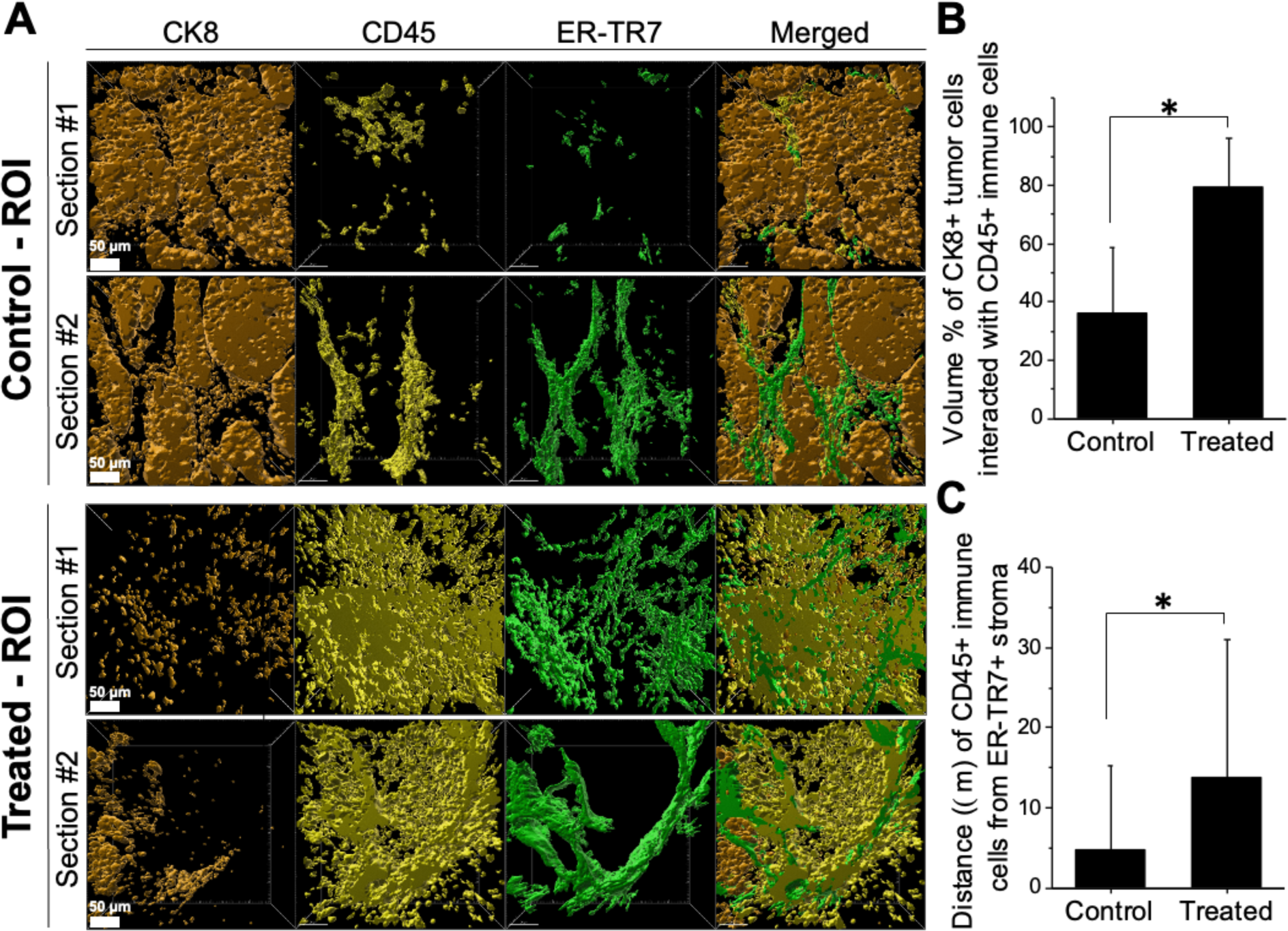
Quantitative cellular resolution 3D spatial analysis of immune infiltration in tumors. **(A)** Segmentation of individual cell type markers in the 3D images of the ROIs from control and DMXAA-treated tumor macrosections. Scale bar: 50 μm. (**B**) Volume percentages (%) of CK8+ tumor cell segments in contact with CD45+ immune cell segments within the 3D images of the ROIs in the control and treated tumors (n=4 per group). *\*P* < 0.05. (**C**) Mean 3D penetration distance (µm) of CD45+ immune cell segments from ER-TR7+ stroma to tumor lesions within the 3D images of ROIs in the control and treated tumors (n=4 per group). *\*P* < 0.05.

### Detection of metastatic lesion in mouse brain using the correlative multiscale 3D imaging method

To extend the application of our method for direct localization of metastatic lesions within normal organs and tissues, we prepared a breast cancer brain metastasis model by intracardiac injection of mouse mammary cancer cells into a mouse. The whole brain tissue was collected at 21 days post-injection and fixed in 4% PFA at 4°C for more than 7 days. Following the established workflow, the brain tissue was optically cleared in organic solvents and scanned by LSFM with a 639 nm excitation laser and corresponding emission filter. The autofluorescence tile images were stitched, and the reconstructed whole mouse brain image was rendered in 3D (**Figure 7A**). Virtual cross-section viewing of the brain image allowed us to readily determine a ROI showing the abnormal anatomical structure in the left cerebral area (**Figure 7B**). Using our ROI tracking methods, we acquired the target 400 μm-thick brain macrosection, labeled it with DAPI, DyLight488-anti-CK8, DyLight550-CD45, and DyLight680-anti-ER-TR7 antibodies, and performed multiplex CLSM imaging after optical clearing in D-fructose solutions. After identifying the selected ROI in the tissue-wide 3D CLSM image of the brain macrosection (**Figure 7C**), we localized CK8+ metastatic cancer cells by cellular resolution 3D CLSM imaging of the ROI (**Figure 7D**). The high-resolution 3D image also showed numerous CD45+ immune cells in the ROI, possibly indicative of an early immune response to metastatic cancer cells in mouse brain^34,35^. This result shows that our correlative multiscale 3D imaging method facilitates identifying and visualizing the *in situ* cellular composition of metastatic lesions and inflammation within large volumetric normal mouse organs and tissues.

**Figure 7.**
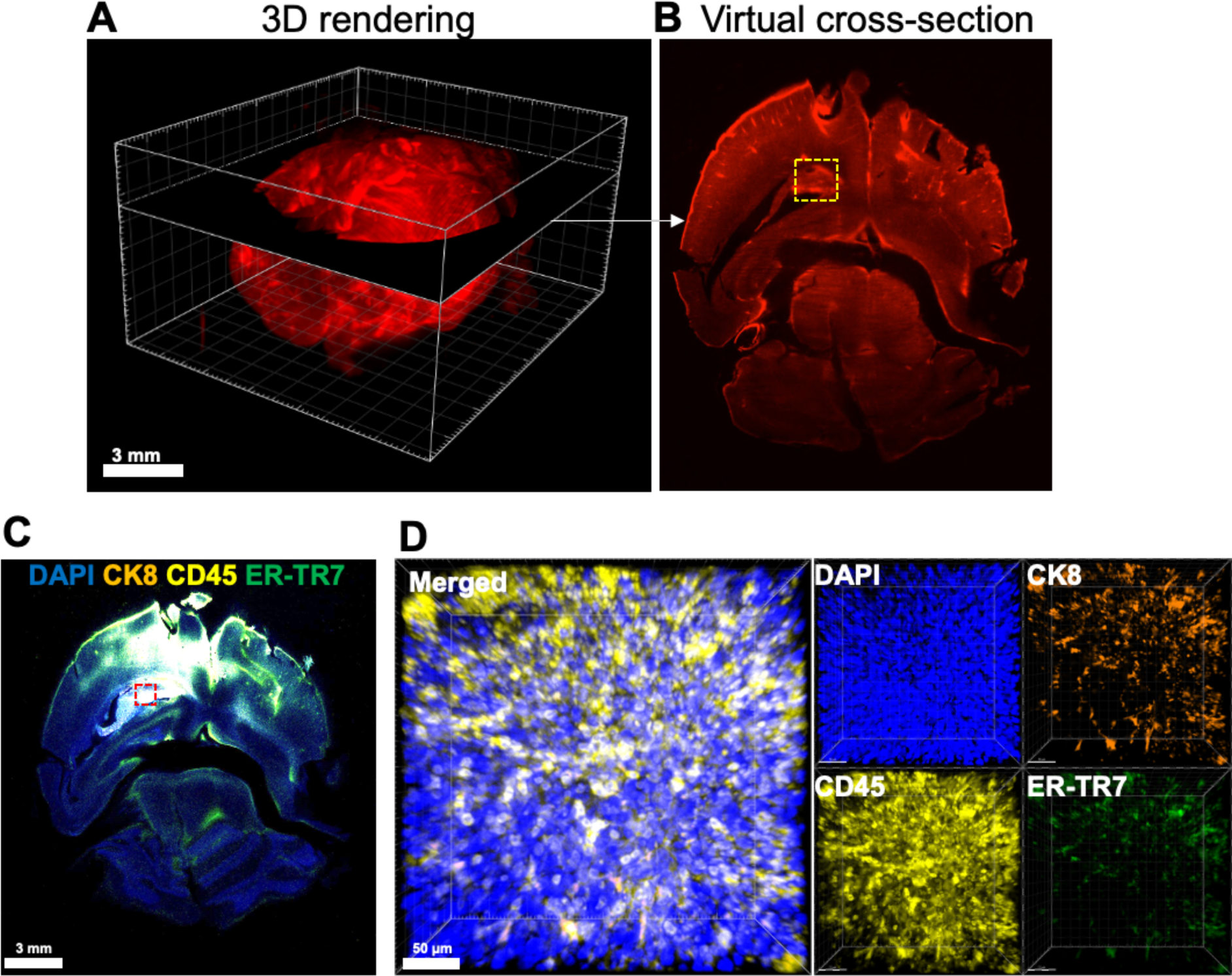
Correlative multiscale 3D imaging to localize a metastatic lesion in a whole mouse brain. **(A)** 3D rendering and virtual slicing of the whole mouse brain images acquired by 3D LSFM scanning of autofluorescence signal. Scale bar: 3 mm. **(B)** Virtual cross-section view featuring a ROI. A yellow dotted box indicates the ROI. **(C)** 3D rendering of the whole brain macrosection image acquired by CLSM. The brain macrosection was stained for DAPI+ (blue) cell nuclei, CK8+ (orange) tumor, CD45+ (yellow) immune cells, and ER-TR7+ (green) fibroblasts. A red dotted box presents the ROI. Scale bar: 3 mm. (**D**) 3D rendering of the ROI obtained by high-resolution CLSM. This cellular resolution 3D image shows the metastatic lesion of CK8+ mouse mammary cancer cells (orange) and the response of CD45+ immune cells (yellow) to the cancer invasion. Scale bar: 50 μm.

## Discussion

Sequential microscopy using two or more different methods enables correlative multiscale imaging of a single biological sample^36,37^. For instance, fluorescence and electron microscopy can be applied sequentially to image the same cell samples^38^. In this process, optical IF microscopy localizes specific intracellular molecules and organelles at subcellular resolution, while subsequent electron microscopy visualizes detailed structures and locations of the target molecules and organelles at nanometer-scale resolution. The multiscale image data can be integrated into a single composite image, providing comprehensive spatial information for cell and molecular biology. Importantly, this correlative multiscale imaging is enabled by a specialized cell sampling technique on an electron microscope grid, which permits the identification of the same single cell object among many cells in both microscopy processes^39^.

To date, a variety of optical microscopy techniques, including LSFM (or ultramicroscopy) and CLSM, have been utilized for 3D imaging of optically cleared tissue samples^40–44^. Each imaging technique offers distinct spatial resolution and FOVs, producing characteristic 3D tissue image data. However, despite the benefits of each optical microscopy technique for 3D tissue imaging, the combination of these imaging techniques for correlative multiscale 3D imaging of tissues has remained largely unexplored. This is primarily due to the absence of a method that enables the assignment of ROI positions within volumetric tissue samples and tracking them during sequential optical microscopy processes.

LSFM is widely used together with organic solvent-mediated optical tissue clearing methods for imaging large tissue samples, such as whole mouse brains and tumors^45–47^. Light sheet scanning with a low-magnification objective (1x or 4x) enables the rapid capture of autofluorescence images with a large field of view, which is useful for defining ROIs within large 3D tissues. However, these low-resolution LSFM images generally provide limited information about the cellular and molecular compositions of the selected ROIs. To address this challenge, efforts are underway to develop methods that allow sequential analysis of cells and molecules in the ROIs of optically cleared tissues defined by LSFM imaging. The Ertürk group recently introduced two post-LSFM ROI analysis approaches^48^. The first method, called DISCO-mass spectrometry (MS)^48^, used LSFM imaging to define ROIs within optically cleared whole mouse brains. After reversing the tissue clearing protocol, the brain tissue is frozen and cryo-sectioned to a thickness of 12 µm. The tissue sections are then subjected to laser capture microdissection (LCM) for spatially selective collection of tissue samples from the ROIs and subsequent proteomic analysis. Although the report demonstrated the feasibility of the method, challenges remain in identifying ROI-containing tissue sections from thousands of cryosections of the whole mouse brain and in registering LCM-mediated tissue section images with the whole brain image from LSFM. The second method, DISCO-bot^48^, utilized a newly developed robotic arm installed in a light sheet microscope system to directly extract tissue samples from ROIs within an optically cleared whole mouse body. A needle on the top of the robotic arm is inserted into the mouse body during real-time LSFM imaging, and negative-pressure aspiration permits the collection of tissue and cell samples from the ROIs. This method enables proteomic profiling of the samples from defined ROIs within whole mouse bodies and whole human hearts using MS analysis. However, this invasive approach disrupts the structural integrity of the tissue in the ROIs, making the method incompatible with investigations of spatial cell organization and cell-cell interactions within the tissue environment. Moreover, the requirement for the specialized instrument, ‘LSFM-bot’, also limits the accessibility of this method to broader research fields.

Our group has previously demonstrated that the switching of optical tissue clearing process from organic solvent (i.e., DBE and ECi) to water-based solution (i.e., D-fructose) allows for multiscale 3D imaging of mouse lung tissues by sequential LSFM and CLSM^24^. However, the different spatial resolution image data could not be correlated to each other due to the loss of the positional information of ROIs after tissue macrosectioning. To overcome this challenge, here we developed a simple but robust method for tracking ROIs in large-sized tissue samples, such as a whole mouse tumor and brain, by employing a UVA dye. Spiropyran transitions from a closed spiropyran form (SP, transparent) to an open merocyanine form (MC, purple) upon exposure to UV light. Under visible light, however, it reverses^49^. This unique optical feature of spiropyran allows for the visible marking of the positions of ROIs on the agarose gel surrounding a mouse tumor or brain using a UV light sheet beam in light sheet microscope system. The additional physical marking preserves the positional information of the ROIs during the subsequent reversal of the organic solvent clearing process and macrosectioning. Multiplex IF and CLSM imaging of the macrosections with high-magnification objectives ultimately provide 3D visualization of various cell types within the ROIs. The LSFM and CLSM images were readily registered together to generate correlative ‘whole mouse tumor (or brain)-to-cellular’ resolution 3D images (**Supplementary Video 1**). With these 3D image data, we performed quantitative analysis of structural remodeling and immune infiltration in the TME of a mouse mammary TUBO tumor to evaluate the effects of DMXAA treatment at multiscale resolutions. By assessing correlated multiscale image data from a single tumor tissue, we can provide integrated spatial information on the heterogeneity of the TME and the immune response to therapy. This approach suggests the potential utility of our method for evaluating cancer therapies from a multiscale spatial perspective. Furthermore, our method enabled direct localization of a metastatic lesion within a whole mouse brain as well as the local metastatic cancer cells and immune cell response. This result demonstrates that our method has potential as a useful imaging tool for detecting and analyzing metastatic lesions within large-sized tissues in tumor metastasis research.

Our ROI tracking method and its associated workflow have proven to enable correlative multiscale 3D optical microscopy of mouse tumors and other normal tissues. However, we recognize the need for further improvements to make the method more user-friendly and capable of gathering more spatial biology information from tissue samples. The first improvement required is the development of a one-direction or slow-reversible UV-activated (UVA) dye. Due to the relatively fast reversion of spiropyran to its deactivated form after exposure to UV light sheet, immediate physical marking is necessary before the visible ‘purple’ line on the agarose gel disappears. If there are multiple ROIs in a tissue sample, it becomes necessary to repeatedly withdraw and return the tissue sample from the LSFM chamber to physically mark each ROI. The second improvement involves extending our imaging workflow to 3D super-resolution optical microscopy (e.g. structured illumination (SIM) microscopy^50^). If we develop a method that allows for marking and tracking the X-Y position of a target single cell within a ROI of a tissue macrosection, correlative ‘whole mouse tumor (or organ)-to-subcellular resolution’ 3D imaging will become feasible. The final improvement needed is to adapt our recent high-power LED-mediated 3D photobleaching technique to enhance multiplexing in the imaging of diverse cell types within tissue macrosections^51^. This adaptation will enable comprehensive cellular profiling of ROIs. We anticipate that these further improvements will enable robust, correlative multiscale and multiplex 3D imaging of large volumetric tumors and other tissues. This advanced 3D optical microscopy method will be useful across a broad range of research fields, including cancer and inflammatory diseases, as well as in the development of new effective therapies.

## Methods

### Mouse mammary tumor model and treatment with DMXAA

TUBO cells, derived from a spontaneous mammary tumor in a BALB-NeuT female mouse^52^, were obtained from Dr. Stephen Kron (University of Chicago) and cultured in Dulbecco’s Modified Eagle’s Medium-High glucose (Millipore Sigma, Cat#: D5671) supplemented with 20% heat-inactivated FBS, and 1% penicillin in 5% CO_2_ and at 37°C. BALB/c female mice (6-8 weeks old) were purchased from Envigo. 2×10^5^ TUBO cells in 50 µl PBS were injected into the right 4^th^ mammary fat pad of a BALB/c female mouse. TUBO tumors were developed and reached 5-8 mm in diameter at 10-14 days after injection of the cancer cells. DMXAA (10 μL at 1 mg/mL in PBS; Millipore Sigma, Cat#: D5817) was directly injected into a TUBO tumor in the mouse using a 29G x 1/2” insulin syringe under inhalational anesthesia using isoflurane. At 2 days post-treatment, the mouse was sacrificed, and the treated tumor was collected. An untreated TUBO tumor was also collected as a control.

### Mouse model of breast cancer brain metastasis

Luciferase-expressing E0771 cells, derived from a spontaneous mammary tumor in a C57BL/6 female mouse, were cultured in RPMI media with 10% FBS, 1% L-Glutamine and 1% penicillin in 5% CO_2_ and at 37°C^53^. C57BL/6J female mice (5-7 weeks old) were purchased from Jackson Laboratories (Stock No: 000664). Following the established protocol^54^, a mouse was anesthetized with ketamine/xylazine and 1×10^5^ E0771 cells were injected in the left ventricle of the heart in 100 µl PBS using a 26G x ½” syringe. The mouse was scarified and the whole brain tissue was harvested at 21 days post-injection of cancer cells.

### Fluorescent antibodies

Primary monoclonal antibodies were purchased from the commercial venders: anti-CK8 (BioLegend, 1E8), anti-CD45 (BioLegend, 30-F11) and anti-ER-TR7 (BioXcell, ER-TR7). Antibodies at a concentration of 0.5 mg/ml in PBS (pH8.0) reacted with amine-reactive fluorophores, including DyLight488-NHS ester (Thermo Scientific, Cat#: 46402), DyLight550-NHS ester (Thermo Scientific, Cat#: 62262) and DyLight680-NHS ester (Thermo Scientific, Cat#46418), at a molar ratio of 1:20 (antibody:fluorophore). Unconjugated, free fluorophore molecules were removed by dialysis purification using cassettes (MWCO 10K) in 1 L PBS at 4°C for 3 days with daily PBS refreshment. Fluorescent antibody solutions were stored at 4°C with protection from light until used.

### Tissue processing and organic solvent-mediated optical tissue clearing

The collected tumor tissues were fixed in 2% paraformaldehyde (PFA) solution (prepared with PBS) for 30 mins at room temperature. The mouse brain tissue was fixed in 4% PFA solution for more than 7 days at 4 °C. After fixation, the tumor and brain tissues were washed with cold PBS several times. The tissues were then dehydrated through sequential incubation in 20%, 50%, and 100% (v/v) ethanol (EtOH) solutions, prepared with deionized (DI) water. The pH of these EtOH solutions were adjusted to 7.8 using triethylamine (Millipore Sigma, Cat#471283) before being applied to the tissue samples. Afterwards, the dehydrated tissues were embedded in individual 2% agarose gel (LE Quick Dissolve Agarose, GeneMate) in a 24 or 12-well plate. The tissues in agarose plugs were further incubated in 100% (v/v) EtOH (pH 7.8) two or three times for complete dehydration. Dibenzyl ether (DBE) (Millipore Sigma, Cat#108014) and ethyl cinnamate (ECi) (Millipore Sigma, Cat#112372) solvents were used as clearing agents for reflective index (RI) matching. Before the clearing process, the pH of both DBE and ECi solutions were adjusted to 7.8 with triethylamine. The tissues were immersed in DBE containing 0.3% (v/v) 1-thioglycerol as an antioxidant (Millipore Sigma, Cat#471283) for 1 day with 3 or 4 solvent refreshments. Subsequently, the tissues were transferred to ECi containing 0.3% (v/v) 1-thioglycerol for 2 days with 2 or 3 solvent refreshments. Tissue dehydration and optical clearing processes were performed using air-tight glass vials covered with aluminum foil. All processing steps were conducted at 4°C under gentle agitation, except for ECi-mediated clearing, which was carried out at room temperature.

### 3D light sheet fluorescent microscopy (LSFM) of optically cleared tissues

The optically cleared mouse tissues were imaged using a light sheet fluorescence microscope (Ultramicroscope Super Plan, Miltenyi Biotec) equipped with a 5.5 Megapixel sCMOS camera (Andor Neo) and a MI Plan 4× DC57 objective (NA: 0.35) with a dipping cap. The individual tissues, embedded in agarose plugs, were mounted on a flat sample holder using super glue, transferred to the designated mounting frame, and placed into the sample chamber of the light sheet microscope, filled with ECi solvent. During mounting, the top edge of the agarose gel was slightly trimmed to mark the X-Y orientations of the tissue samples. Whole mouse tumor and brain tissues were scanned using a single-sided light sheet beam source (sheet NA: 0.141, thickness: 3.4 µm, width: 40%) at defined pixel sizes (X×Y×Z = 2.71 × 2.71 × 1.7 µm). A 639 nm excitation laser and a 665-695 nm emission filter were employed, with the laser set to 100% transmission power, to detect tissue autofluorescence. The tile image data were transferred to a PC workstation, and 3D images of the entire tissues were reconstructed using Imaris Stitcher software (version 10.0.0). With 3D rendering and virtual section viewing of the tissue images in Imaris, the ROIs within the tissues were defined.

### Optical and physical marking of ROIs

Spiropyran (or NitroBIPS, Millipore Sigma, Cat# 273619) was dissolved in ECi (pH 7.8) at a concentration of 10 mg/ml as a stock solution, using a hot air gun to warm the solution during dissolution. The tissue, embedded in an agarose plug attached to the sample holder, was removed from the chamber and placed in a glass vial containing 10 mL of ECi (pH 7.8) with 0.5 mL of the spiropyran stock solution. After incubating for 8-12 hours at room temperature, the tissue was returned to the ECi-filled sample chamber of the light sheet microscope. The X-Y orientation of the tissue was matched to the previous position by referring to the marker on the top edge of the agarose gel. Since the flat plastic sample holder remained attached to the bottom of the tissue, it could easily be placed on the mounting frame and returned to the chamber without significant changes in the sample’s Z-axis position for repeat LSFM imaging. This setup facilitated the iterative identification of defined ROI positions in the tissue through live scanning in LSFM. After positioning the light sheet beam at the Z-axis corresponding to the first ROI, the tissue was exposed to a single-sided UV light sheet beam (405 nm) for 3 mins under the same microscope setup as for 3D LSFM imaging. The tissue was immediately pulled out, and the surface of the agarose gel surrounding the tissue was cut along a purple-colored line using a razor blade. The process of ‘scanning, positioning, and optical and physical marking’ was repeated to mark additional ROIs.

### Reversing the clearing process and macrosectioning

After physical ROI marking, the tissues were rehydrated by reversing the organic solvent-based tissue clearing process as follows: the tissues were incubated in a descending gradient of EtOH (prepared in DI water) - 100% at room temperature, 50%, 20%, and finally PBS at 4°C. Several additional washes with fresh PBS followed to ensure complete rehydration. The 400 μm-thick macrosections retaining the ROIs were obtained by vibratome sectioning of the rehydrated tissues, guided by the physical marks on the agarose gels. The macrosections were stained with 0.7 µL of DAPI (5 µg/mL, Thermo Scientific, Cat#: 62248) and 5 µL of each fluorescently labeled anti-CK8, CD45, and ER-TR7 antibody solution (0.5 mg/mL) in 0.5 mL of staining buffer composed of RPMI1640 cell culture media (GibcoTM, Cat#: 22400089) with 1% (w/v) IgG-free bovine serum albumin (BSA, Millipore Sigma, Cat#: A2058) for 18 hours at 4°C under gentle shaking. After washing three times in cold PBS, the macrosections were optically cleared by incubation in a series of 10 mL D-fructose (Millipore Sigma, Cat#: F0127) solutions at varying concentrations (20%, 50%, 80%, and 100% (w/v) in 10 mM phosphate buffer (pH 7.8)), each for 30 mins at 4°C under gentle agitation.

### 3D confocal laser scanning microscopy (CLSM) of tissue macrosections and ROIs

High-resolution 3D multiplex IF images were acquired using an upright confocal fluorescence microscope (Caliber ID, RS-G4) equipped with a 20x/0.8 NA dry objective (Olympus, UPLXAPO20X) and a 40x/1.4 NA oil immersion objective (Olympus, UPLXAPO40XO). 3D CLSM imaging was performed at defined voxel sizes (X×Y×Z) for the whole tissue macrosection (0.614 × 0.614 × 3.05 µm³) and cellular resolution 3D ROI images (0.313 × 0.313 × 1.52 µm³) with 8 frame averaging. The imaging utilized the following excitation and emission wavelengths: a 405 nm excitation laser with a 415-485 nm emission filter for DAPI; a 488 nm excitation laser with 506-594 nm emission filter for DyLight488; a 561 nm excitation laser with 574-626 nm emission filter for DyLight550; and a 640 nm excitation laser with 662-738 nm emission filter for DyLight680.

### Testing optical and physical ROI marking methods using a fluorescent microbead

A single fluorescent microbead, 200 µm in diameter (Cospheric, Cat# 120725-100-1), was placed at the center of a 2% agarose plug before the gel solidified. The agarose plug then underwent the organic solvent-based tissue clearing process as previously described. The microbead within the agarose plug was detected by LSFM using a 4× immersion objective, a 488 nm excitation laser, and a 500-550 nm emission filter. After incubation in a spiropyran solution, the agarose plug was exposed to a UV light sheet beam at the Z-axis position where the microbead was detected for optical ROI marking. Subsequent physical ROI marking was performed using a razor blade. The agarose plug was then cut into 400 µm thick serial sections with the guidance of the physical ROI mark. The gel macrosection containing the physical mark was imaged using CLSM to confirm the presence of the single microbead, using a 10x air objective, a 488 nm excitation laser, and an open emission filter.

### 3D image processing and quantitative analysis

3D images of tumor and brain tissues acquired by LSFM and CLSM were reconstructed using Imaris Stitcher (version 10.0.0). 3D rendering and virtual cross-section viewing of these images were performed with Imaris software (version 10.0.1). To determine the similarity between the virtual tumor cross-section images selected from 3D LSFM and CLSM images (**Figure 3**), the ‘BigWarp’ plugin in Fiji software was utilized to align and register the selected virtual cross-section images without image deformation. Each image was segmented based on its autofluorescence signals and converted into 8-bit binary file using Fiji. We then masked the tumor areas in the images and executed the ‘Clean Outside’ command. The outer layer of each segmented tumor image was obtained by processing in Fiji software: first by implementing ‘Process -> Binary -> Fill Holes’ for the segmented image, followed by ‘Process -> Binary -> Outline.’ The similarities between LSFM and CLSM images were quantitatively determined using the ‘SSIM Index’ plugin in Fiji software. The SSIM ratio for the positive control was determined using identical LSFM or CLSM images. For the negative control, the SSIM ratio was calculated by cross-comparing LSFM and CLSM whole cross-section images between different control and treated tumor groups. For instance, we compared the LSFM image of control section #1 with the CLSM image of treated section #1, and the LSFM image of control section #2 with the CLSM image of treated section #2. To measure the volume percentage (%) of each cell type within the control and DMXAA-treated tumor macrosections (**Figure 5B**), we first performed hyperstack segmentation of DAPI+, CK8+, CD45+, and ER-TR7+ cell areas in the 3D whole tumor macrosection images using individual cutoff thresholds in Fiji software. In this process, all pixels in a raw channel image corresponded to a specific cell or nucleus marker. Then, we converted the raw 3D images into 8-bit binary hyperstack images, measured the ‘area’ value of each segmented cell type marker in every stack of images, and calculated the ‘total volume’ by multiplying the sum of these ‘area’ values by the Z-interval (3.05 µm). The volume percentage (%) of each cell type was determined by *V %* = (Total volume of a cell type / Total volume of DAPI+ tumor parenchyma) × 100. For measuring the distribution patterns of CK8+ and CD45+ cells across the whole macrosections (**Figure 5C**), the segmented hyperstack CK8+ and CD45+ macrosection images were displayed in a ‘Z-project’ view with the ‘sum slices’ mode in Fiji software. The intensity profiles for CK8+ and CD45+ areas along a straight line across the individual macrosection images were obtained using the ‘plot profile’ command in Fiji software.

For quantitative 3D spatial analysis of the ROIs within the tumor macrosections (**Figure 6**), we segmented CK8+, CD45+ and ER-TR7+ cells in high-resolution hyperstack images using the ‘Surface’ modules in Imaris software. The cutoff thresholds were determined by interactive comparison of each cell mark signal in the raw image data. We used ‘Split touching Objects (Region Growing)’ module for accurate and fully representative individual cell surface segmentation. We further processed the segmented images with ‘Overlapped Volume’ filter as inclusive or exclusive criteria for quantitative determination of CK8+ cancer - CD45+ immune cell interaction (**Figure 6B**). The segmented images were also processed with ‘Shortest distance to surface=ER-TR7’ to measure the 3D penetration distance of CD45+ immune cells from ER-TR7+ stroma to CK8+ tumor (**Figure 6C**).

We processed and registered the multiscale LSFM and CLSM images of the control tumor into a 3D image using both Fiji and Imaris software. Initially, we identified the image stacks in the LSFM hyperstack autofluorescence image of the whole tumor that corresponded to the CLSM image stacks of two selected IF-labeled tumor macrosections. This process was guided by the recorded Z-axis positions (i.e., the numbers of LSFM slice image files) of the LSFM whole tumor image, which were exposed to a UV light sheet beam for optical marking of the ROIs. Next, we used the ‘BigWarp’ plugin in Fiji software to align the two CLSM macrosection images with the corresponding LSFM stack images and exported the processed macrosection images as separate image files. Additionally, using the ‘BigWarp’ plugin in Fiji software, we aligned the cellular resolution 3D IF image to fit the correct ROI position within the macrosection image, based on its recorded location information (position #2) on the macrosection #2, and exported the processed image as a new file. After converting all the image files to Imaris format, we combined them into a single Imaris file using the ‘Add Channel’ feature. The correlated multiscale 3D image of the control tumor and its ‘Google Earth’-like view stream video were created in Imaris software (**Supplementary Video 1**).

## Statistical analysis

Values are displayed as mean ± standard deviation (SD), and statistics were run using unpaired t-tests in Prism software. If the *P* value is less than 0.05, it is considered “significant.”

## Supporting information

Googlemapviewdata

## Acknowledgements

We thank Drs. Stephen Kron and Vytautas Bindokas at the University of Chicago for providing the TUBO cell line and consulting on image processing and analysis.

## Ethics Approval and Consent to participate

Animal experiments were approved by the Animal Care and Use Committee of the University of Illinois Chicago and in compliance with the Guide for the Care and Use of Laboratory Animals as adopted by the U.S. National Institutes of Health

## Author contributions Statement

J.Z. and S.S.-Y.L. conceived the concept and designed the study. Y.-C.W., X.C., P.P., E.E.E, Z.Z., and J.Z. prepared, treated and processed mouse tumor tissues. J.Z. performed immunofluorescence, and LSFM and CLSM microscopy. X.C. and J.Z. conducted 3D image data processing and quantitative analyses. J.Z. and S.S.-Y.L. wrote and edited the manuscript. All authors approved the final manuscript.

## Data Availability Statement

All data collected from this study are available in this published article and supporting information.

## Supplementary Figure Legends

**Supplementary Figure 1.**
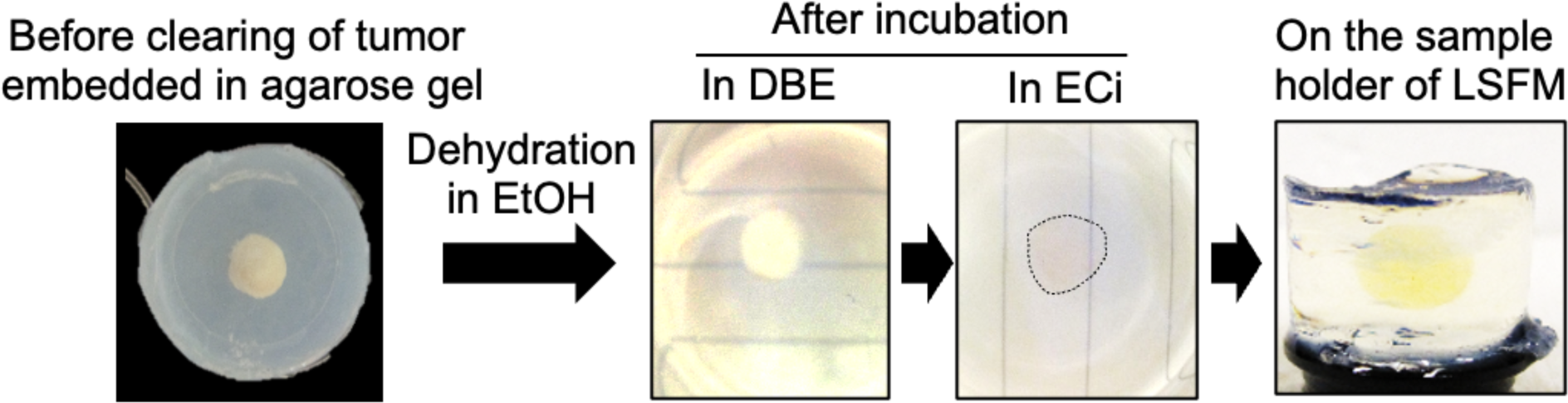
Optical tissue clearing process of tumors using organic solvents. After embedding in agarose gel, the mouse tumor was dehydrated in EtOH, and cleared through sequential incubations in DBE and ECi. The cleared tumor within agarose gel was mounted on a flat sample holder using super glue.

**Supplementary Figure 2.**
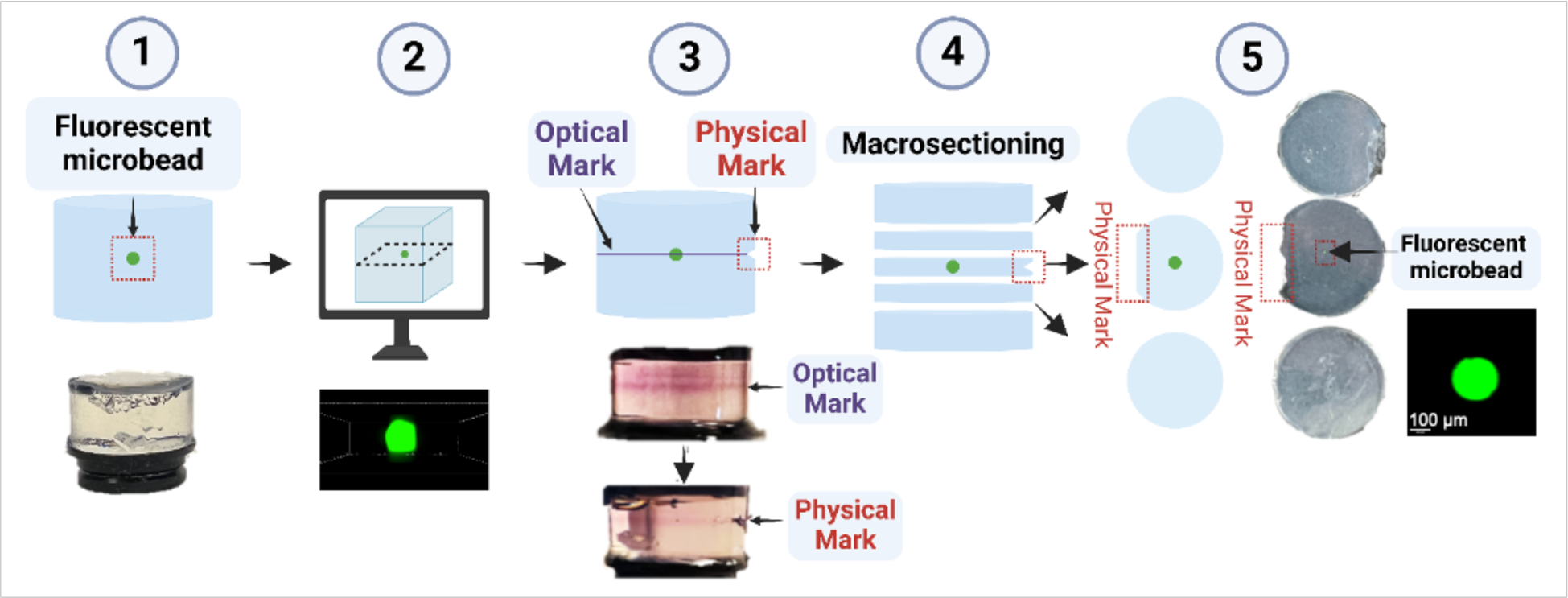
Examination of optical and physical ROI marking procedures using a fluorescent microbead. **(1)** A single fluorescent microbead was placed in the center of an agarose plug to create an artificial ROI. The bottom photograph shows the agarose plug containing the microbead. **(2)** After the organic solvent-mediated clearing process, 3D LSFM imaging localized the microbead within the agarose plug. The bottom LSFM image displays the microbead. **(3)** Using spiropyran, a UV light sheet beam, and a razor blade, the Z-position of the microbead in the agarose plug was marked optically and then physically. The bottom photographs show the agarose plug with optical and physical ROI marks. (**4**) Serial macrosectioning of the agarose plug was performed guided by the physical ROI mark. (**5**) The left-side illustration and right-side photographs of 400 µm-thick agarose macrosections demonstrate the presence of the physical mark on the macrosection, aiding in the identification of the macrosection containing the microbead. The right bottom CLSM image localized the microbead in the macrosection with the physical mark. Scale bar: 100 μm.

**Supplementary Figure 3.**
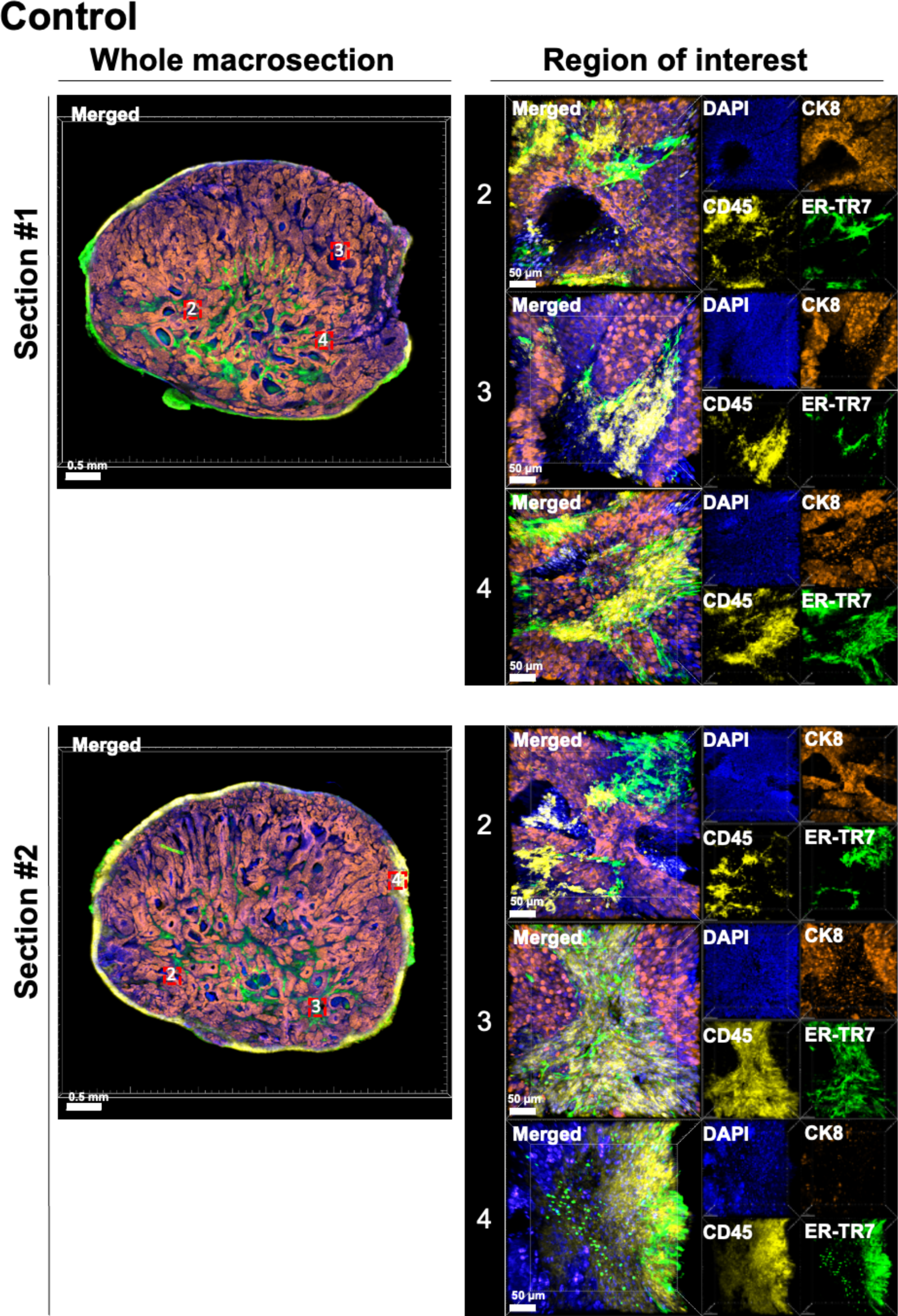
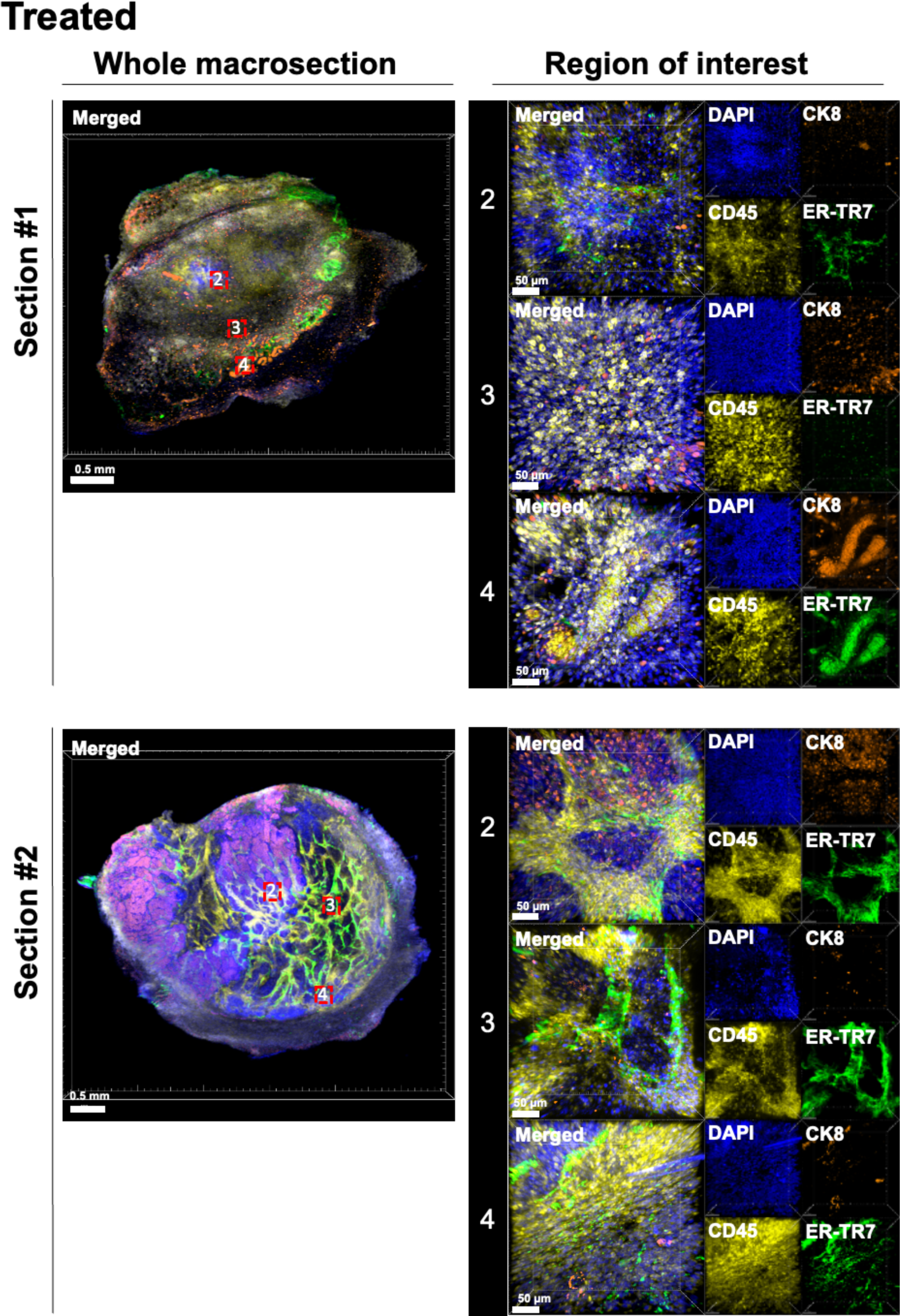
High-resolution 3D multiplex images of other ROIs in the control and DMXAA-treated tumor macrosections. The tumor macrosections were stained for DAPI+ (blue) cell nuclei, CK8+ (orange) tumor, CD45+ (yellow) immune cells, and ER-TR7+ (green) fibroblasts. Red dotted boxes with numbers on the left-side whole macrosection images indicate individual ROIs (#2-#4). The corresponding high-resolution 3D CLSM images of the ROIs are displayed on the right side. Scale bar: 0.5 mm (left) and 50 µm (right).

---

**Supplementary Video 1. 3D rendering of the registered multiscale 3D image of the control mouse tumor**

